# Structure of the ForT/PRPP complex uncovers the mechanism of C-C bond formation in C-nucleotide antibiotic biosynthesis

**DOI:** 10.1101/2020.03.26.009662

**Authors:** Sisi Gao, Ashish Radadiya, Wenbo Li, Huanting Liu, Wen Zhu, Valérie de Crécy-Lagard, Nigel G. J. Richards, James H. Naismith

**Affiliations:** Research Complex at Harwell, Didcot, OX11 0FA, United Kingdom; BSRC, University of St. Andrews, St. Andrews, KY16 9ST, United Kingdom; School of Chemistry, Cardiff University, Cardiff, CF10 3AT, United Kingdom; Division of Structural Biology, University of Oxford, Oxford, OX3 7BN, United Kingdom; Department of Chemistry and California Institute for Quantitative Biosciences, University of California, Berkeley, CA 94720, USA; Department of Microbiology, University of Florida, Gainesville, FL 32611, USA; Foundation for Applied Molecular Evolution, Alachua, FL 32415, USA; The Rosalind Franklin Institute, Didcot, OX11 0FA, United Kingdom

**Keywords:** C-nucleosides, Formycin, X-ray Structure, Docking, Enzyme Mechanism

## Abstract

C-C bond formation is at the heart of anabolism and organic chemistry, but relatively few enzymatic strategies for catalyzing this reaction are known. The enzyme ForT catalyzes C-C bond formation between 5’-phosphoribosyl-1’-pyrophosphate (PRPP) and 4-amino-*1H*-pyrazole-3,5-dicarboxylate to make a key intermediate in the biosynthesis of the C-nucleotide formycin A 5’-phosphate; we now report the 2.5 Å resolution structure of the ForT/PRPP complex and thus locate the active site. Site-directed mutagenesis has identified those residues critical for PRPP recognition and catalysis. Structural conservation with GHMP kinases suggests that stabilization of the negatively charged pyrophosphate leaving group is crucial for catalysis in ForT. A mechanism for this new class of C-C bond forming enzymes is proposed.

**Entry for the Table of Contents:** 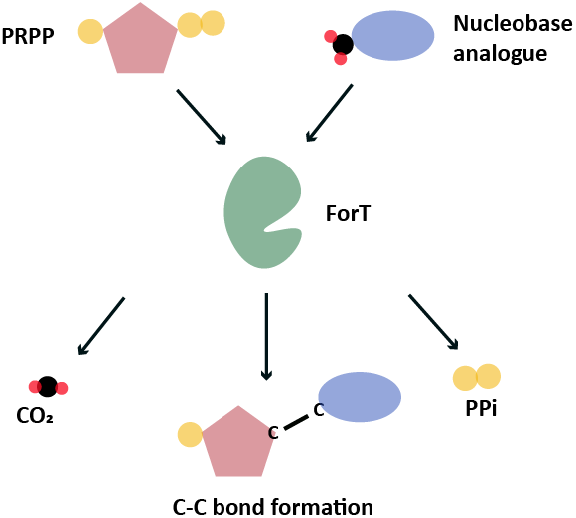

A new class of enzymes catalyse C-C bond formation by irreversible CO_2_ and pyrophosphate production.

---

There is renewed interest in the synthesis and characterization of C-nucleosides^[1]^ driven by the potency of GS-5734 against ebola,^[2]^ and coronaviruses, including those that cause SARS^[3]^ and MERS.^[4]^ In C-nucleosides, the nucleobase is connected to C-1’ of the sugar ring by a C-C bond rather than a C-N bond, thereby rendering the molecule stable and altering its stereoelectronic properties.^[5,6]^ With the exception of pseudouridine synthase,^[7,8]^ relatively little is known about the structures and catalytic mechanisms of enzymes that form C-ribosides and C-glycosides.^[9,10]^ Recent work identifying the biosynthetic gene clusters for formycin A **1**,^[11–13]^ pyrazomycin **2** (also known as pyrazofurin)^[14,15]^ and showdomycin **3**^[16]^ (Figure 1a) has laid the foundation for obtaining an enhanced understanding of two new enzymes (ForT and PyfQ) that catalyse C-C bond-forming steps in C-nucleoside biosynthesis. Both ForT and PyfQ were originally functionally assigned based on homology to (4-(β-D-ribofuranosyl)hydroxybenzene (RHP) synthase, which mediates a key step in methanopterin biosynthesis.^[17–19]^ These enzymes all utilize 5’-phosphoribosyl-1’-pyrophosphate (PRPP) and an aromatic carboxylic acid to make the new C-C bond (Figure 1b). The liberation of inorganic pyrophosphate and irreversible CO_2_ release provide the driving force for these reactions. Sequence alignments show that these three enzymes are homologous with, and thus likely are evolutionarily related to, homoserine kinase,^[20]^ a member of the GHMP kinase superfamily (Figure S1, Supporting Information).^[21]^

**Figure 1.**
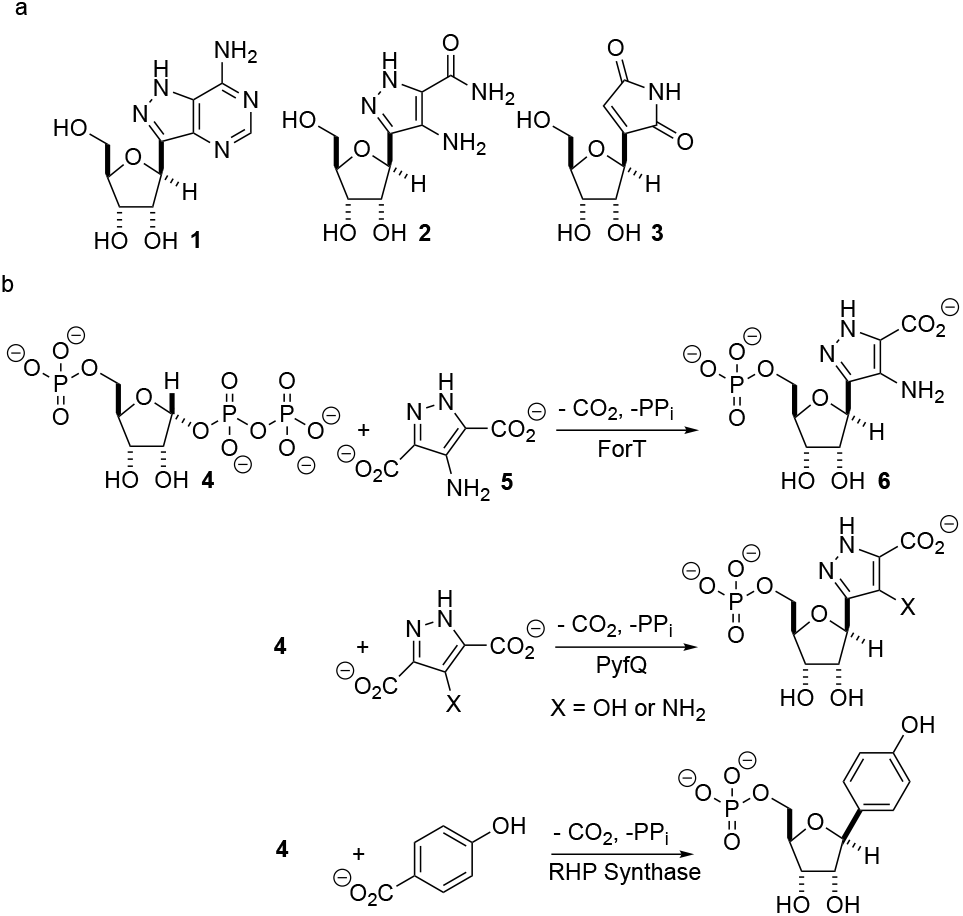
C-nucleotides. (a) Formycin **1**, pyrazomycin **2** and showdomycin **3**. (b) A new class of C-C bond forming enzymes that utilise 5’-phosphoribosyl-1’-pyrophosphate **4** and aromatic carboxylic acids to make C-nucleotides.

Two previous studies^[12,14]^ have shown that the substrate of ForT is 4-amino-*1H*-pyrazole-3,5-dicarboxylate (ADPA) **5** (Figure 1b) and that the enzyme-catalyzed reaction yields C-nucleotide **6** (Figure 1b), an intermediate in formycin biosynthesis. Nothing is known, however, about the structure of ForT or the active site residues that play a role in substrate binding and/or catalysis. We therefore overexpressed and purified recombinant, wild type (WT) ForT in *Escherichia coli*, and grew crystals of the enzyme in the presence of 10 mM PRPP **4**. The resulting crystals were then soaked in 200 mM of PRPP **4** before data collection at the Diamond Light Source, which allowed us to solve the structure of the ForT/PRPP complex to 2.5 Å resolution (Figure 2a and Table S1, Supporting Information). The crystal asymmetric unit contains a ForT monomer with residues 11-171, 180-205 and 209-341 (C-terminus) experimentally located in electron density; we assume that the missing residues are located in conformationally disordered regions. The monomer has three antiparallel β-sheets which form the core of the structure. Two of the sheets share the same elongated strand (residues 11 to 17 for sheet 1; 18 to 24 for sheet 2) and sit end to end (Figure 2a). The third sheet sits opposite, and partially stacks against, sheet 2. There are two α-helices packed against one face of sheet 1 with a third packing against the other two. Sheet 2 has a very small α-helix packed against one of its faces whilst sheet 3 has two helices attached. A bundle of four helices is packed against the ends of sheet 2 and sheet 3. Gel filtration and multi-angle light scattering suggests the enzyme to be a dimer in solution (Figure S2, Supporting Information), and the 2-fold rotation axis in the crystal does result in one (and only one) plausible dimer that relies on contacts between the helical bundles of each monomer (Figure 2b). Interestingly, analysis of this dimeric arrangement with the PISA server^[22]^ assigns the dimer with low confidence (0.3 on a 0 to 1 scale) due to relatively few interactions between the monomers and the limited amount of buried surface area.

**Figure 2.**
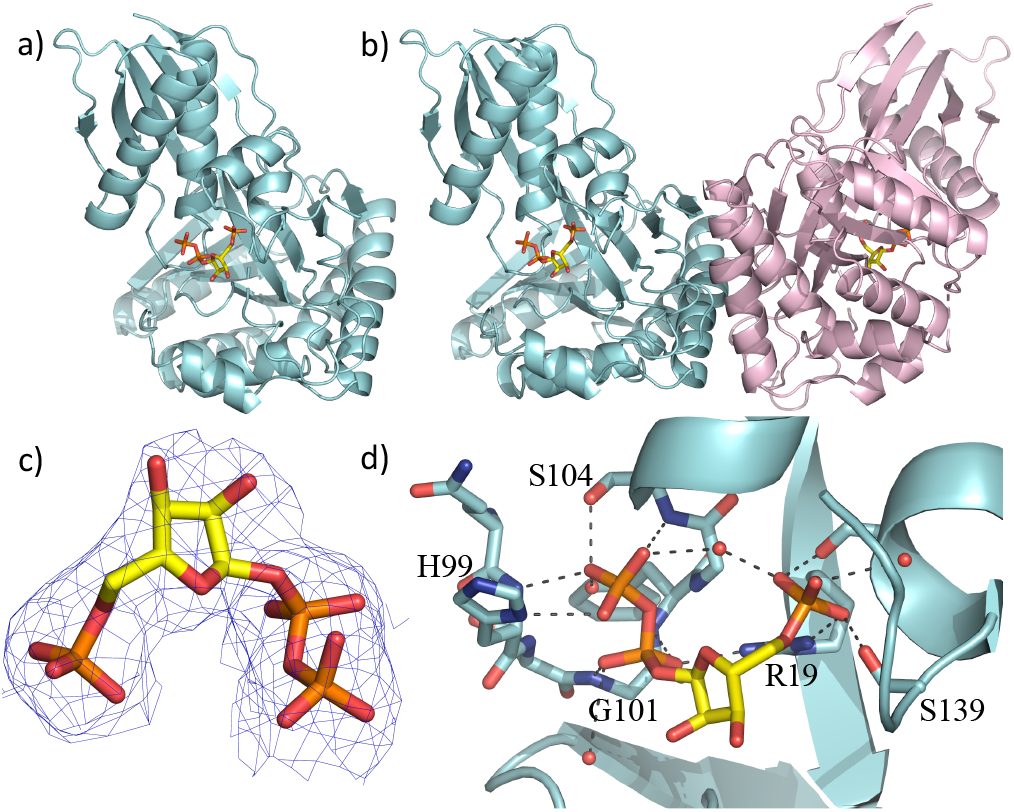
X-ray crystal structure of the WT ForT/PRPP complex. (a) The ForT monomer shown as a cyan cartoon, PRPP, shown in sticks (carbon yellow, phosphorous orange, oxygen red) is at the centre of the monomer. (b) The ForT dimer is generated by crystal symmetry, the second monomer is pale pink cartoon, PRPP is remote from the dimer interface. (c) The final 2Fo-Fc map contoured at 1.2σ for PRPP (shown as Figure 2a) (the original Fo-Fc map is shown in Figure S3, Supporting Information). (d) PRPP is bound to the enzyme by an extensive array of hydrogen bonds. The loop (Gln-98 to Ser-108), characteristic of GHMP kinase superfamily, plays a crucial role in substrate binding. Protein carbon atoms are colored in cyan, nitrogen in blue, other atoms as in Figure 2a.

The sequence of ForT is distantly related to homoserine kinase (< 20% identity), a member of the GHMP kinase superfamily, although there are regions of strong sequence conservation. The ForT structure reveals that the closest structural analogue is indeed homoserine kinase from *Methanocaldococcus jannaschii* (1fwl)^[20]^ (root mean square deviation (rmsd) of 2.6 Å over 266 Cα atoms). Superposition reveals the conservation of the three antiparallel β-sheets and the helices (Figure S3a, Supporting Information). There is a notable difference between the structures centred on the bundle of four helices, ForT has two longer loops (residues 25 to 35 and residues 157 to 181). Homoserine kinase is a dimer and like ForT, the dimer interface uses the same bundle of helices. However, the longer loops in ForT preclude exactly the same dimeric arrangement seen for homoserine kinase; the second monomer in the ForT dimer is rotated by approximately 60° relative to second monomer in the homoserine kinase dimer (Figure S3b, Supporting Information). ForT is also structurally related to mevalonate kinase^[23]^ (PDB 6mde; rmsd 2.6 Å over 262 Cα atoms). The mevalonate kinase has a completely different dimeric arrangement, however, even though it too uses the same helical bundle to form the dimer (Figure S3c, Supporting Information).

The difference electron density map indicated the presence of bound PRPP in the enzyme (Figure S3d, Supporting Information). Pyrophosphate and phosphate only were therefore added to the model and the structure further refined. The resulting difference density connecting the phosphate and pyrophosphate moieties improved allowing fitting of the ribose ring, thereby locating PRPP in the ForT/PRPP complex (Figure 2c). PRPP is located in the centre of the monomer, remote from the dimer interface (Figure 2b,2d). The 5’-phosphate of PRPP sits in a pocket and forms salt bridges with the side chains of Arg-19 and Arg-135; it also interacts with the N-terminus of short helix in the helical bundle. Hydrogen bonds are also observed between the 5’-phosphate and the side chains of Thr-106, Ser-139, Ser-142 and the backbone NH of Ser-142 (Figure 2d). The pyrophosphate moiety sits in different pocket and also makes a salt bridge with Arg-19. The pyrophosphate is hydrogen bonded to all seven backbone amides in a tight turn comprising residues His-99 to Ser-104 as well as with the side chain of His-99 and two water molecules (Figure 2d). This (pyro)phosphate binding tight turn is a conserved feature of the GHMP kinases (Figures 2d and S1, Supporting Information). A bound water also appears to bridge the phosphate and pyrophosphate moieties. Our assignment relies on the observed coordination sphere being consistent with water and not of a magnesium ion. The ribose ring only makes one van der Waals contact with the side chain of Val-294. The weaker density of the ribose also suggests some conformational flexibility. We note that two disordered loops (172 to 179 and 210 to 340) are plausibly within reach of this region and it is thus possible additional protein/ribose interactions may exist.

Using isothermal titration calorimetry (ITC) the dissociation constant of the ForT/PRPP complex was determined to be 4.0 μM (Table S2, Supporting Information). We did not observe any binding of ribose 5-phosphate. Informed by the crystal structure, we prepared a series of site-specific ForT variants by mutagenesis (Table S3, Supporting Information) and used these variants in ITC measurements (Figure S4a, Supporting Information), which confirmed that Arg-19, Thr-106 and Arg-135 are indeed essential for PRPP binding (Table S2, Supporting Information). Replacing Gly-134 by proline decreased PRPP binding by almost 100-fold, consistent with the idea that the conformation of this loop segment is important in recognition.

With the crystal structure in hand, standard modeling methods were used to construct the disordered loops in ForT (Supporting Information).^[24]^ During turnover, the active site has to accommodate the atoms of pyrophosphate and the reaction product **6**. We thus constructed two tautomers of a probe molecule (**7** and **8**), which combines the molecular features of both substrate and product (Figure 1). These “hybrid” probes (Figure 3a, 3b) were manually placed in the structure by simple superimposition on the experimentally located PRPP. The ADPA moiety of the probe molecule has extensive clashes with the protein (Figure S3e, Supporting Information), indicating either the PRPP molecule or the protein (or both) adjust their position upon catalysis. A large positively charged pocket is close the probe (Figure 3c) which would be well matched for the negatively charged ADPA molecule. This pocket is lined by several polar residues (Figure 3d).

**Figure 3.**
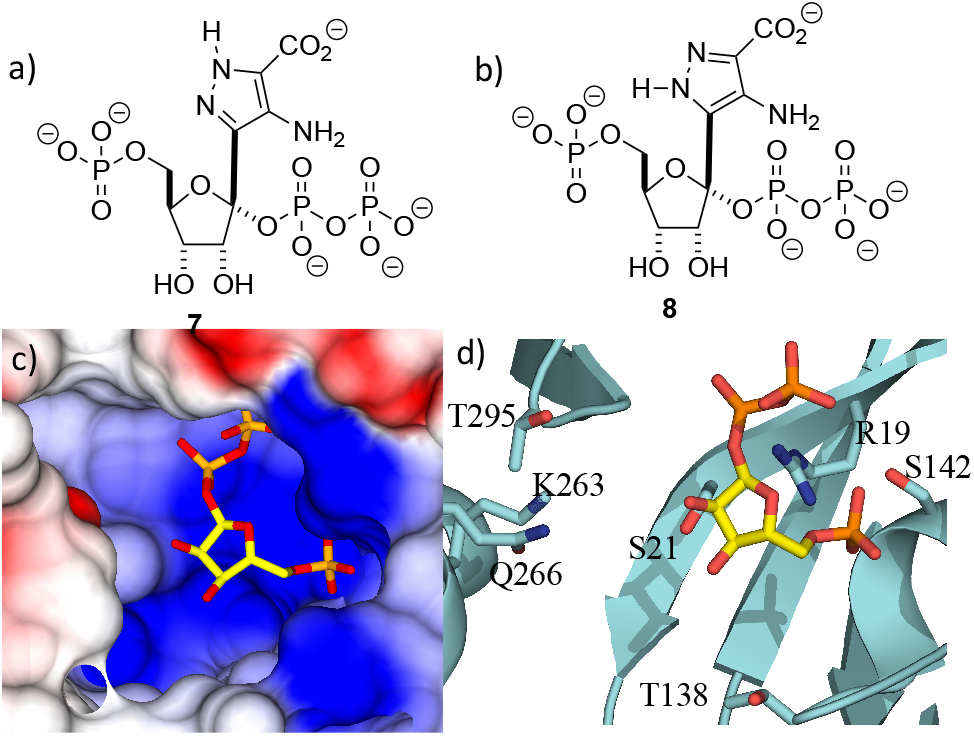
Location of the second substrate (a) Tautomer of a simple model for transition state. (b) Alternative tautomer of model for transition state. (c) PRPP sits a large positively charged pocket that would bind the negatively charged ADPA molecule. The binding pocket is shown as an electrostatic surface. (d)The active site pocket is lined by several polar residues, coloured as in Figure 2d.

We also evaluated the catalytic activity of the recombinant WT ForT used to obtain the X-ray crystal structure. Thus, LCMS confirmed formation of the expected product^[12]^ when the enzyme was incubated with PRPP **4** and ADPA **5**, analysis with LCMS confirmed the production of the expected product^[12]^ (Figure S4b, Supporting Information). The activity of WT ForT and a number of ForT variants were also assayed using membrane-inlet mass spectrometry (MIMS)^[25–27]^ to measure ForT-catalyzed CO_2_ production (Table S2 and Figure S4c, Supporting Information). These experiments showed that WT ForT has a specific activity of 0.002 U/mg under the reaction conditions used to identify the substrates for the ForT-catalysed reaction by Liu and co-workers.^12^ The addition of EDTA to the reaction mixture abolished activity, suggesting that Mg^2+^ is needed for catalytic activity. The ease of the MIMS-based assay also permitted a preliminary kinetic assessment of selected ForT variants (Table S2, Supporting Information). Unfortunately, many of these ForT variants exhibited poor stability under our standard reaction conditions.

It is perhaps surprising that ForT should share structural homology with kinases because phosphorylation and C-nucleoside bond formation are very different chemical transformations. Superposition of the ForT/PRPP complex with that of homoserine kinase bound to phosphoaminophosphonic acid-adenylate ester (AMP-PNP) (PDB 1h72),^[28]^ however, shows that the γ-phosphate of ATP overlaps with the C-1’ phosphate of the pyrophosphate (proximal phosphate) of PRPP (Figure 4a). Similarly, superposing the ForT/PRPP complex with homoserine kinase bound to ADP (PDB 1fwk)^[20]^ shows that the β-phosphate of ADP overlaps with the distal phosphate of the pyrophosphate in PRPP (Figure S3f, Supporting Information). We have previously noted that structural and chemical similarity between an adenylating enzyme and a (non-GHMP superfamily) kinase arose from the shared need to stabilize a negatively charged phosphate transition state^[29]^. In GHMP kinases, the γ-phosphate undergoes nucleophilic attack and the enzyme, using the GHMP kinase loop and a Mg^2+^ ion^[20]^, stabilizes the developing negative charge (Figure 4b). Both the γ-phosphate in homoserine kinase and the proximal phosphate of PRRP in ForT make very similar interactions with the GHMP kinase loop in their respective structures. This degree of conservation we take to imply that in both enzymes, stabilization of the negatively charged phosphate is key. In ForT, this need for stabilization implies that the C-1’–pyrophosphate bond undergoes significant dissociation in the rate determining step forming an S_N_1-like transition state with an oxocarbenium ion (Figure 4c). The breakage of the pyrophosphate ribose bond would permit the movement of the ribose ring which we noted earlier, is required for formation of the adduct with ADPA. The oxocarbenium ion would be highly reactive to the electron rich aromatic ADPA molecule and the subsequent elimination of CO_2_ would act as an irreversible step. Based on the similar to homoserine kinase, we presume the apparent requirement for Mg^2+^ arises from stabilisation of pyrophosphate.

**Figure 4.**
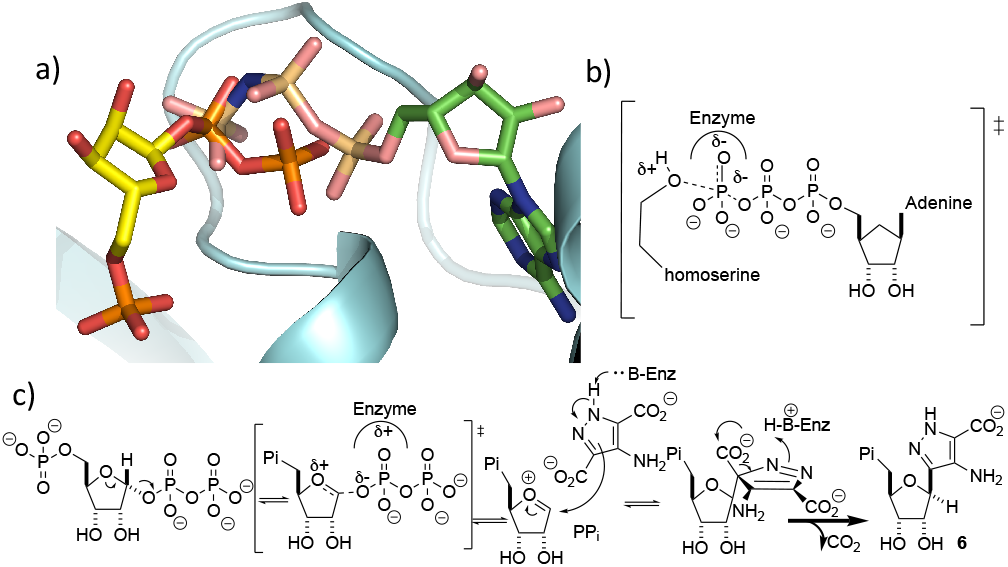
Mechanistic implications of the ForT/PRPP structure. (a) Overlay of PRPP in the ForT/PRPP complex with AMP-PNP homoserine kinase complex. The proximal pyrophosphate of PRPP overlaps with the γ-phosphate of AMP-PNP. (b) The transition state for γ-phosphoryl transfer in the reaction catalysed by homoserine kinase. (c) Proposed mechanism for ForT-catalysed C-C bond formation.

C-nucleotides, such as formycin, are promising medicines for the treatment of viral diseases. As part of our studies of this pathway ^[30]^, we now report the structure of ForT, the first for of a novel class of C-C bond forming enzymes. We have identified key residues that mediate substrate binding and catalysis. These findings suggest a mechanism for C-C bond formation that provides new insights into the different ways enzymes can utilize the high energy PRPP molecule.^[31,32]^

## Acknowledgements

We thank for the Diamond Light Source for beam time allocation and beam line staff for assistance with data collection. Funding for these studies was provided by BBSRC (BB/T006161/1 & BB/T006188/1 to J.H.N. & N.G.J.R., respectively).

